# Serum proteomics reveals high-affinity and convergent antibodies by tracking SARS-CoV2 hybrid immunity to emerging variants of concern

**DOI:** 10.1101/2024.10.02.616394

**Authors:** Anand Patel, Thiago Lima, Richard Carson, Qiulong Huang, Stefano R. Bonissone, Natalie Castellana

## Abstract

The rapid spread of SARS-CoV2 and continuing impact on human health has prompted the need for effective and rapid development of monoclonal antibody therapeutics. In this study, we interrogate polyclonal antibodies in serum and B cells from whole blood of three donors with SARS-CoV2 immunity to find high-affinity anti-SARS-CoV2 antibodies to escape variants. Serum IgG antibodies were selected by affinity to the receptor-binding domain (RBD) and non-RBD sites on the spike protein of Omicron subvariant B.1.1.529 from each donor. Antibodies were analyzed by bottom-up mass spectrometry, and matched to single- and bulk-cell sequenced repertoires for each donor. Antibodies observed in serum were recombinantly expressed, and characterized to assess domain binding, cross-reactivity between different variants, and capacity to inhibit RBD binding to host protein. Donors infected with early Omicron subvariants had serum antibodies with subnanomolar affinity to RBD that show binding activity to a newer Omicron subvariant BQ.1.1. The donors also showed a convergent immune response. Serum antibodies and other single- and bulk-cell sequences were similar to publicly reported anti-SARS-CoV-2 antibodies, and characterized serum antibodies had the same variant-binding and neutralization profiles as their reported public sequence. The serum antibodies analyzed were a subset of anti-SARS-CoV2 antibodies in the B cell repertoire, which demonstrates significant dynamics between the B cells and circulating antibodies in peripheral blood.

## 1 Introduction

Severe acute respiratory syndrome coronavirus 2 (SARS-CoV-2), the virus that causes coronavirus disease (COVID-19), continues to be a serious global health threat as new escape variants emerge. Early in the COVID-19 pandemic, researchers sought to develop high-affinity therapeutic monoclonal antibodies (mAbs) to rapidly clear SARS-CoV-2 from infected patients, and prevent onward transmission in high-risk environments [7]. In late 2021, a number of antibodies (e.g., bamlanivimab, casirivimab, sotrovimab, bebtelovimab, and tixagevimab) received emergency use authorization from the United States Food and Drug Administration (FDA) and European Medicines Agency (EMA) for early infection and treatment of mild to moderate disease. However, with the global spread of SARS-CoV-2, new variants emerged with enhancing mutations that escape immunity. Due to ineffectiveness against the currently dominant Omicron subvariants, the antibodies that were developed against the wild-type strain are no longer recommended for therapeutic use. In March 2024, Pemivibart was approved for the prevention of COVID-19 for moderate-to-severely immune compromised individuals. Derived from an existing mAb, adintervimab, that was developed to target the Delta subvariant, Pemivibart is capable of neutralizing the JN.1 Omicron subvariant. While vaccine programs have significantly reduced lethality of the disease, there is still a need for surveillance of new escape variants, and rapid discovery and development of therapeutics to treat infection.

Severe acute COVID is not the only health concern. An estimated 5%-30% of individuals who have been infected with the virus have Long COVID with persistent or recurrent symptoms, such as fatigue, weakness, and brain fog, lasting four weeks to years. The etiology of Long COVID is poorly understood, with potentially multiple mechanisms contributing to disease onset [8], including persistence of dormant SARS-CoV-2 that initiates disease recurrence, or uncoordinated humoral and cellular immune responses. Similar to severe acute COVID [21], Long Covid may require a coordinated anti-SARS-CoV-2-specific T and B cell response for protection from disease [29]. There are conflicting reports as to whether elevated anti-SARS-CoV-2 antibody titers or slow neutralizing antibody response to infection are correlated with Long COVID [29, 23, 5, 17]. A deeper characterization of antibodies in serum may provide clearer insights into the causes of Long COVID.

The humoral immune response has been primarily investigated by measuring antibody titers against various antigens, and more recently with single-cell technologies that sequence Ig transcripts from individual B cells (scBCR-seq). Measuring titers provides an overview of abundance and/or strength of antibody binding, but is incapable of distinguishing if the strength is from one antibody clone, many related clones, or many distinct clones. On the other hand, single-cell technologies provide a highly detailed view of the cellular component, showing diverse paired heavy and light chain sequences for thousands of cells, some showing clonal expansions. Many research groups have applied the single-cell approach to identify neutralizing antibodies by screening B-cell receptors (BCRs) from peripheral blood of convalescent or vaccinated individuals for activity to viral antigens [12, 30, 18]. To initiate infection, coronaviruses enter a host cell by relying on the engagement of the receptor-binding domains (RBD) of spike proteins on the viral envelope with angiotension-converting enzyme-2 (ACE2) receptors on epithelial cells. Both RBD and spike protein have been used as antigens to screen B cells for producers of highly potent therapeutic antibodies [18]. However, the B cells accessible via peripheral blood are only a sub-sampling of the millions of naive and memory B cells that are present in bone marrow. Serum antibodies, on the other hand, specifically antigen-specific antibodies observed in the same peripheral blood, represent a functional state of the humoral immune response with a highly selected and focused set of clones [13]. Deep characterization of antibodies is now possible with proteogenomics methods that combine next-generation sequencing of Ig transcripts and mass spectrometry analysis of antibody proteins [6, 14, 2, 15]. The discordance can be stark with *<*0.1% of peripheral B cell clones represented by serum antibodies at steady-state [14].

Broad adoption of proteogenomic methods has been challenging due to required resources and expertise across genomics, proteomics, and bioinformatics [4]. Despite the challenges, proteogenomics has been applied to more deeply characterize anti-SARS-CoV-2 serum antibodies. In one example, antibodies from four study subjects with mild COVID and varied virus-neutralization titers were further characterized by proteogenomics to show that 84% of clones in serum are directed to non-RBD epitopes of the spike protein [27]. Additional efforts have confirmed the effects of immunological imprinting, where initial exposure to a virus or vaccine directs the type of immune response upon subsequent exposures [26]. Individuals infected with wild-type SARS-CoV-2 are imprinted with an serum antibody response directed at non-RBD spike protein epitopes, whereas the response for vaccinated individuals tends to primarily be targeted against RBD epitopes [28].

We used proteogenomics in this study to profile serum antibodies from two donors who were vaccinated and later developed an Omicron breakthrough infection, and one donor who was naturally infected with SARS-CoV-2 and later vaccinated to investigate how imprinting may provide protection against escape variants. All three donors experienced mild symptoms after vaccination and infection, and whole blood was collected at least 4 months from the most recent boost or breakthrough infection. We compared the serum antibodies to the B cell repertoire generated by single cell sorting and sequencing, revealing the presence of class-switched clones between the B cell compartment and secreted antibodies. Many of the cellular BCRs and serum antibodies observed showed sequence similarity to previous publicly reported anti-SARS-CoV2 antibodies. We selected a panel of antibodies for recombinant expression and characterization, which revealed epitope coverage of both the receptor binding domain (RBD) and N-terminal domain of the spike protein. We screened the antibodies for binding to the SARS-CoV-2 wild-type strain and two Omicron subvariants: B.1.1.529 and BQ.1.1. We found monoclonal antibodies retaining binding activity to a variant-of-concern BQ.1.1, despite reports of the variant first appearing more than five months after each donor’s infection.

## 2 Materials and Methods

### 2.1 Sample Collection and Processing

Samples were collected from three donors who had been vaccinated and infected with SARS-CoV-2. Whole blood was drawn by venipuncture from each donor, between 120-161 days after exposure. 150mL to 200mL of blood was collected from each donor, and stored in Vacutainer tubes with EDTA (BD #366643). The study design was approved and materials were collected following IRB protocol #22-ABTE-101, and reviewed by Pearl Pathways, LLC.

## 2.2 B cell enrichment and paired VH:VL sequencing

On the same day as collection, B cells were enriched using RosetteSep Human B Cell Cocktail (StemCell #15064). The SepMate protocol was followed using Lymphoprep density gradient medium, followed by dilution with Dulbecco’s PBS with 2% FBS (StemCell #07905) as medium in SepMate-50 tubes. Following separation by centrifugation, the top layer containing serum antibodies was extracted by serological pipette into a PETG media bottle, and the second layer contain B cells was poured out into separate collection tubes for further washing with PBS. Cells were counted by Countess 3, and then pelleted and cryopreserved in CryoStor CS10 (# 07930) buffer.

B cells were thawed at 37C and washed twice with EasySep buffer (PBS, 2% fetal bovine serum, 1mM EDTA; StemCell). Cells were incubated with 3ug of biotinylated omicron spike trimer (Acro Biosystems # SPN-C82Ee) for 15 min before proceeding with the isolation (StemCell Human Biotin Positive Selection Kit #17663). Isolated cells were resuspended in EasySep buffer and loaded onto a 10X Chromium Next Gem Chip K. BCR libraries were constructed using the Chromium Next GEM Single Cell 5’ v2 V(D)J kit on a 10X Genomics Chromium Controller following user guide protocol [9].

Remaining non-BCR isolated cells were added to 2mL of lysis buffer, and 500uL was used for RNA extraction following Zymo Quick RNA protocol [20]. Extracted RNA was quantified by spectrophotometry (VarioSkan uDrop Duo plate). Bulk IgG transcript libraries were constructed using SMARTer Human BCR IgG IgM H/K/L Profiling Kit (Takara #634466).

### 2.3 Ig and BCR transcript sequencing and repertoire construction

scBCR-seq data was generated on the Illumina HiSeq platform in 2 × 150bp paired-end read format generating 380M reads, *≈*128M per donor. Data was analyzed using CellRanger 7.0.1 in VDJ mode. The consensus sequences in fasta file format were then annotated using an in-house pipeline and stored in AIRR-seq format [25].

Bulk Ig transcript sequencing data was generated on the Illumina NextSeq 1000 platform in 2 × 300bp paired-end read format. Transcript sequencing data was processed starting from FASTQ files with an in-house pipeline for trimming adaptors, pair-stitching, and filtering of low quality reads. Briefly, raw paired-end reads were filtered to remove pairs with either read containing an N base or mean quality value lower than 20. Reads were stitched together using prefix-suffix pairwise alignment, and the maximum quality value in the overlap portion was retained. Stitched reads were then error corrected using an in-house variant of a Hamming graph approach, similar to [22]. All reads in a dense subgraph of the Hamming graph are error corrected by consensus, with the new clustered read ascribed abundance equal to the number of raw reads in the cluster. The cluster size is used as the RNA abundance value for the error corrected sequence. Any clusters with abundance less than two are removed as they likely contain errors and cannot be corrected.

BCR and Ig transcript repertoire sequences were V(D)J annotated using alignment [3] to human functional V, D, and J gene references from IMGT [16]. Complementarity-determining regions (CDRs) and somatic hypermutation events are inferred from gene alignments. An AIRR-seq file containing constructed repertoire, V(D)J labeling, CDR identification, and mutation calling was generated for each sample. AIRR-seq files were concatenated to use as a single database in searching mass spectra to estimate false discovery rates. Transcripts that share the same CDR3 are referred to as clonal. The identified CDR3s are delimited by the residue after the conserved second cysteine and residue preceding the J region tryptophan or phenylalanine prior to conserved WGXG or FGXGT motif depending on heavy or light chain.

### 2.4 IgG and antigen-enrichment from plasma

IgG antibody was purified from 10mL of plasma from each donor by affinity purification with Protein G. Briefly, plasma was diluted to 20mL in PBS buffer, and passed through a 0.2 micron filter. Filtrate was than loaded onto a 5mL HiTrap Protein G HP column (Cytiva #17040501), washed with 25mL of PBS, and eluted with 15mL of DEA at pH 11. HiTrap column was regenerated with washes of 30mL DEA, and 30mL PBS, and re-used for each sample.

IgG antibody was fractionated for affinity to SARS-CoV2 Omicron (B.1.1.529) RBD (Acro #SPD-C522e) or (Acro #SPN-C5224) spike trimer. Affinity resin for purification was prepared by conjugating 0.5mL NHS agarose resin with 300ug of RBD or spike at 4C overnight. Resin was split into three columns, one for each donor.

Two sets of purifications were performed using 6mL and 10mL of IgG from each donor. For the first set, 6mL of IgG from each donor was loaded onto RBD columns, and incubated for 30min at room temperature (RT). The flow-through was collected and the column was washed with 6mL PBS, and eluted with 0.25mL DEA, pH 11 in four fractions, and neutralized with 25uL 1M Tris pH 7. The flow-through was purified one more time to deplete anti-RBD IgGs with their respective columns. Flow-through was then loaded onto columns with coupled spike trimer, and purified as above, except spike+ IgG was eluted in one fraction with 0.75mL DEA, pH 11. For the second purification set, 10mL of IgG from each donor was loaded onto regenerated RBD columns. The RBD-depleted flow-through was then loaded onto spike columns, and eluted in one fraction with 0.75mL DEA, pH 11.

In total, four fractions were analyzed per patient, two 6mL RBD+ IgGs fractions, and 6mL and 10mL RBD-depleted spike+ IgGs. Quantity and purity of IgG was estimated using absorbance at A280 and SDS-PAGE.

### 2.5 Bottom-up mass spectrometry analysis

Antigen-purified IgG from each donor were separated into heavy and light chain by SDS-PAGE, and digested into peptides by four different proteases for tandem mass spectrometry (MS/MS) analysis. Briefly, IgG samples were reduced with NuPAGE Sample Reducing Agent (#NP0009 Thermo), and split into four wells with 5ug-10ug per lane of a 10% Bis-Tris agarose gels (NuPAGE #NP0306BOX Thermo). Gels were run in MOPS buffer, and heavy and light chains were visualized with SimplyBlue SafeStain (#LC6060 Thermo). Heavy chain bands were cut from the gel, and sliced into approximately 1mm squares. Gel pieces were destained, reduced, and alkylated, and normalized to digestion buffers. Briefly, gel pieces were washed with 500uL of 50% acetonitrile (ACN), dried, and resuspended in 300uL of 100mM ammonium bicarbonate (ABC) with 10mM dithiothreitol (DTT). Reduction of disulfide bonds was carried out for 30min at 50C. Iodoacetic acid (IAA) was added for a final concentration of 60mM, and incubation for 30 min at RT in the dark. The buffer containing ABC, DTT, and IAA was then removed, and the gel pieces were washed twice using 50% ACN, with a final dry-down by speed-vac for 15 min. The dried gel pieces were then prepared for digestion by chymotrypsin, elastase, pepsin, and trypsin by resuspending in 300uL of their respective protease buffers: 100mM Tris-HCl 10mM CaCl2, pH 8.0 for chymotrypsin; 50mM Tris-HCl, pH 9.0 for elastase; 100mM ABC, pH 1.5 for pepsin; and 100mM ABC, pH 8.5 for trypsin. 1ug of each protease was added per digest, and the samples were incubated overnight (16-18 hours) at 37C. Proteases were inactivated with 1% formic acid. Peptides were extracted from gel pieces with resuspension in 120uL of 50% ACN, 5% formic acid buffer, incubated for 45min at RT, sonicated for 5min, and transferring extracted peptides to a new glass vials. Extraction was repeated once more, and peptides were dried in a speed-vac, and resuspended in 300uL of 100mM ABC, 0.5% trifluoroacetic acid (TFA). Peptide digests were further desalted with C18 solid-phase extraction (Empore 96-well #6030SD CDS Analytical). Eluted peptide samples were dried-down, and resuspended in 50uL of 2% ACN 0.1% formic acid, incubated for 5 min. at RT, then transferred to mass spec vials for analysis; injection volumes of 10-20 uL (corresponding to 20-40% of the sample digests) were used.

Peptide sample digests were analyzed using an Aurora Ultimate Column (1.7um, C18, 25cm × 75um ID #AUR3-15075C18 IonOpticks) on a Dionex Ultimate 3000 HPLC coupled to an Orbitrap Eclipse mass spectrometer. Buffer A was 2% ACN, 0.1% formic acid in Optima-grade water and Buffer B was 80% ACN 0.1% formic acid in Optima-grade water. Peptides were eluted using the following 2-hour gradient (percent Buffer B) at 0.3 uL/min: 10% to 28% (100 min.), 28% to 55% (20 min.). MS1 scans were performed with a cycle time of 3s at 60K resolution, a scan range of 300-1600 m/z, an automatic gain control (AGC) target of 4e5, and a maximum injection time of 50ms. Precursors for MS2 were selected by passing the monoisotopic precursor selection (MIPS) filter, an intensity threshold of 50K, a charge of +2 to +8, and dynamic exclusion of 60s after two scans. Fragmentation of peptide precursors at an Orbitrap resolution of 30K with quadrupole isolation and an AGC target of 4e5 took place using a hybrid high-energy collision dissociation (HCD)/electron transfer dissociation with supplemental HCD activation (EThcD) method: All peptide precursors were fragmented by HCD, with some of the same precursors undergoing EThcD. HCD fragmentation was done using a quadrupole isolation window of 1.6 m/z, and was stepped at 25%, 35%, and 50% normalized collision energies (NCE); the scan range was set to 101-3000 m/z, and the maximum injection time set at 54 ms. EThcD was performed using a quadrupole isolation window of 3.0 m/z, with calibrated charge-dependent electron transfer dissociation (ETD) parameters and 15% NCE supplemental activation; the scan range was set to 105-3000 m/z and the maximum injection time set at 250 ms.

### 2.6 Clone selection with Alicanto data analysis

Alicanto is a proteogenomic platform [2] for identifying antigen-specific antibodies present in donor serum. Briefly, Alicanto matches tandem mass spectra generated from antigen-purified antibody protein from serum to an antibody repertoire generated from RNA found in B cells. Alicanto scores each antibody clone, defined as unique amino acid CDR3, in the repertoire based on supporting peptide-spectrum match evidence. The scoring model selects clones interpreting fragmentation from each spectra for identifying the entirety of the clone’s CDR3 sequence in the context of similar clones. Only clones with at least 2 unique residues were selected, after masking antibody sequences for repeating 12-mers.

Alicanto was applied to the RBD+ and the spike trimer+ fractions matched with the Ig and BCR repertoire from three donors. Serum clones were identified from the 6 Alicanto runs, and a set of 17 antibodies from BCR repertoires were selected for recombinant expression and characterization. Candidates from all fractions were evaluated including 1 from AB2, 12 from AB5, and 4 from AB6. 12 of the candidates were found in RBD-purified proteomic fractions and 10 were found in spike-trimer+ proteomic fractions.

Repertoire amino acid sequences were identified as public clones by matching heavy chain V-gene annotations and requiring CDR3s to be at least 10 amino acids and align with *≥*80% match to an antibody reported in CoV-SAbDab [19]. For each sequence with a clone match, a single entry in SAbDab was chosen to propagate characterization information by aligning heavy variable chains, and selecting the entry with greatest percent match.

### 2.7 Recombinant antibody expression and purification

The variable region sequences were codon optimized, synthesized with signal peptides, and cloned into an expression vector containing a human IgG1/Kappa for expression in monoclonal format (mAb). A large-scale plasmid preparation was performed, followed by 3mL transient expression in CHO-K1 cells. Supernatant was collected for a one-step purification using Protein G. Quantity and purity was estimated using absorbance at A280 and SDS-PAGE.

### 2.8 ELISAs for domain mapping and neutralization

Expressed mAbs were tested for binding to a panel of targets from three SARS-CoV2 variants. Each antibody was tested for binding to RBD, N-terminal domain (NTD), and spike trimer from wild-type virus, Omicron B.1.1.529, and a newer variant of concern, BQ.1.

MAbs were also tested for neutralization using a SARS-CoV-2 Variant Inhibitor Screening Assay (sVNT; #VANC00, R&D Systems). Briefly, 100uL/well of His-tagged RBD from Omicron B.1.1.529 at 6ng/uL (#SPD-C522e, Acro) was captured on a plate using anti-His capture antibody for 75min at RT. mAbs were normalized to 0.46mg/mL, and incubated on the plate to capture antibody binding to the immobilized RBD. Each mAb was 3-fold diluted serially into six wells with two replicates per dilution. Next, biotinylated ACE2 was added to the plate, and incubated for 90min at RT. Plates were washed four times, followed by detection with streptavidin-HRP and colorimetric substrate. The reduction of absorbance at 450nm with increasing concentrations of mAb indicated ACE2 blocking and were identified as neutralizers in the assay.

### 2.9 Biolayer Inteferometry (BLI)

Expressed antibodies with observe binding activity were selected for biolayer inteferometry (BLI) analysis on an Octet RED96 (Sartorius). Expressed monoclonal antibody was immobilized to the AHC sensor at 5ug/mL concentration. Recombinant His-tagged RBD Omicron B.1.1.529 was used as analyte with varying concentration in 3-fold and 2-fold dilution starting at 1uM and 50nM in PBS pH7.4 buffer, and 0.02% Tween-20. Kinetic analysis was performed with 120s capture, 120s association, and 180s or 600s dissociation. Sensor regeneration was performed with 10mM Glycine pH1.5 with three 5s cycles.

## 3 Results

### 3.1 Serum proteomics detects antibodies to Omicron virus variant months after exposure

B cells from each donor were sorted for cells expressing BCRs that bind to SARS-CoV2 Omicron spike protein, and single-cell sequenced to obtain a repertoire of paired heavy and light V(D)J antibody sequences. The number of distinct heavy chain sequences range from 17,812 to 22,817 in the BCR repertoire (see Figure 2). As expected when sampling the antigen-experienced memory B cells, the number of clones, as defined by unique amino acid CDR3s, is similar to that of distinct sequences in the repertoire. On the other hand, the Ig repertoire, which is derived from bulk B cells and selected for IgG class transcripts, has a lower clone to sequence ratio and 33%-46% of sequences have somatic mutations differences from other sequences of the same clones. The Ig and BCR repertoires have distinct clones, with only 0.5%-2.0% of clones in common between the repertoires. The low overlap reflects that the sequenced repertoires are only a sub-sampling of the total transcriptional repertoire present in peripheral blood.

**Figure 1.**
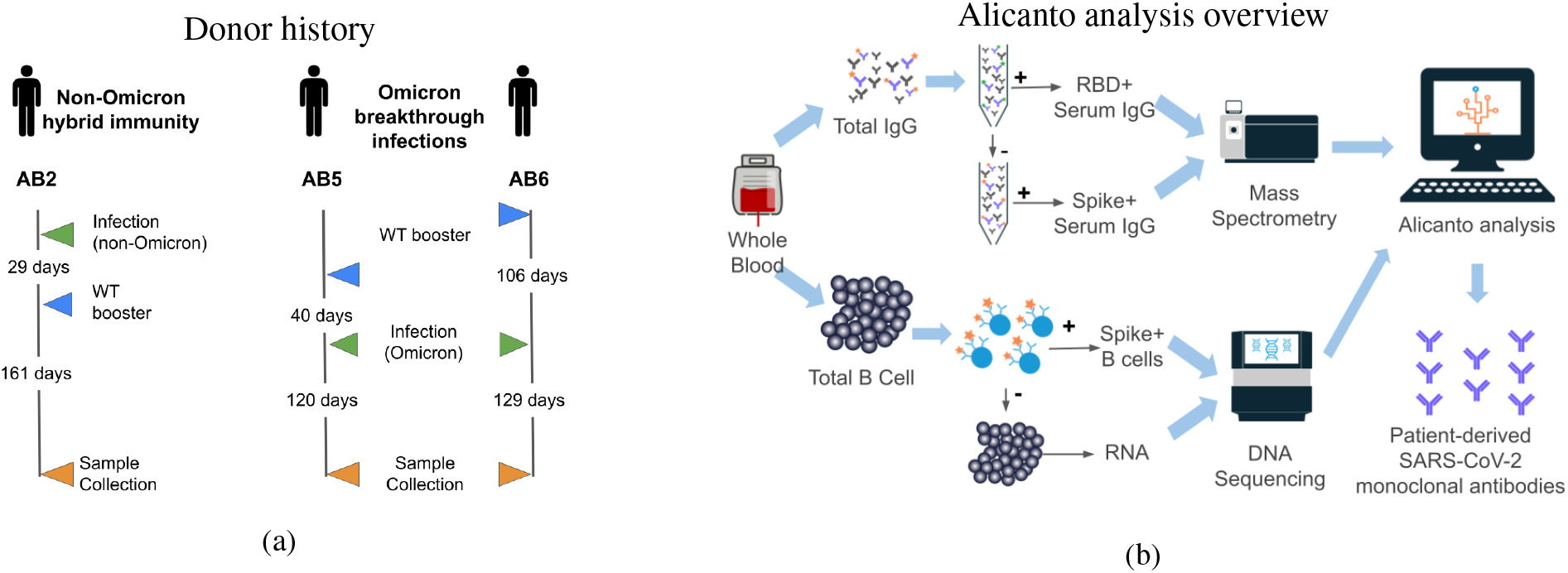
a) Schedule of vaccination and infections for three donors prior to blood collection for this study. Blood was collected between 120-161 days from the most recent exposure. Donor AB2 was only exposed to wild-type vaccines and a non-Omicron natural infection. Donors AB5 and AB6 received wild-type vaccines and suspected Omicron natural infections. b) Overview of antibody and B cell processing, data generation, and analysis. Total B cells and plasma were collected from peripheral blood. RBD- and spike-reactive antibodies were purified from plasma and analyzed by tandem mass spectrometry. Spike-reactive B cells were enriched from total B cells and processed for single B-cell V(D)J transcript sequencing. The Alicanto software analyzes the mass spectrometry data and next-generation sequencing data to identify a set of patient-derived antibody candidates.

**Figure 2.**
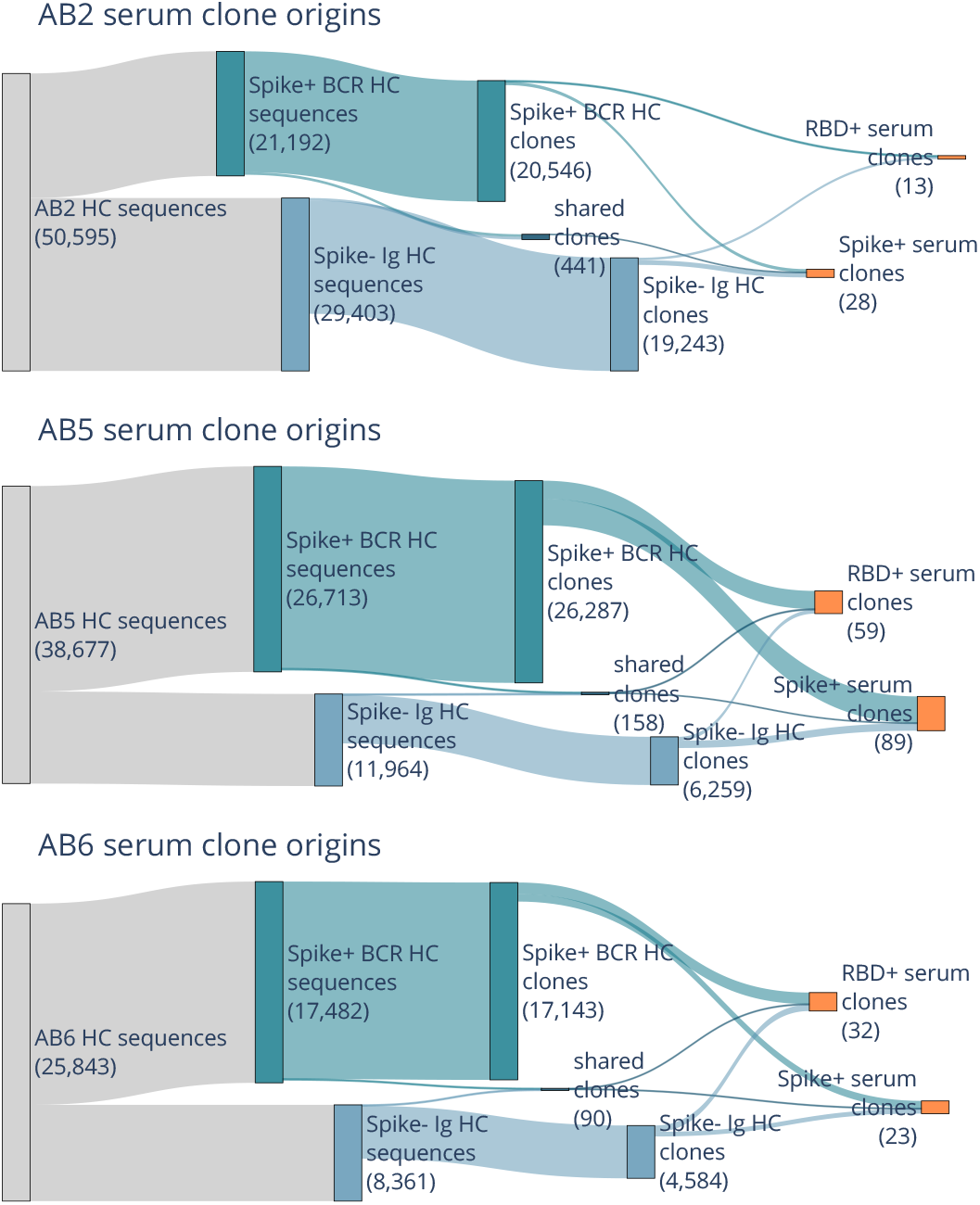
Sankey plots for donors AB2, AB5, and AB6 shows heavy chain BCR (dark blue) and Ig (light blue) repertoire sequence counts collapsing to BCR, Ig, and shared repertoire clones (second blue boxes), and then serum selected anti-RBD and anti-spike clones (orange). Counts of sequences or clones are shown in parenthesis. Few clones are shared between BCR and Ig repertoires, but are still present in serum.

The population of B cells and circulating antibodies in peripheral whole blood reflect the dynamics of the immune response. Even though BCRs and antibodies were both enriched for specificity to spike protein, 99.4%-99.8% of BCR repertoire clones are not observed as circulating IgG antibodies in serum. The highest number of clones in serum are from donors AB5 and AB6, both of whom were suspected to be exposed to an Omicron variant through natural infection. Despite not having being exposed to Omicron antigens, either by vaccination or natural infection, donor AB2 still produced serum antibodies that are reactive to Omicron variant RBD and spike, suggesting AB2 retained some immunity to emerging new variants.

### 3.2 Serum IgG appear in BCR repertoire as non-class switched IgM

The BCR repertoire is primarily dominated by IgM class clones with fewer than 3 amino acid mutations from a germline V-gene, and are likely from naive B cells. While the serum antibodies analyzed are all IgG as a result of Protein G purification prior to antigen purification, they can map to sequences of a different antibody class in the BCR repertoire due to the same B-cell clone having expanded into differentiated and class-switched cells. In the AB5 BCR repertoire, 84% of repertoire clones are IgMs compared to 6% being IgG (see Figure 3). Of the AB5 clones in serum, 67% of spike+ and 47% of RBD+ IgG clones are IgM class in the BCR repertoire, and most of these clones are naive. The Ig repertoire, which is focused on B cells expressing IgG class transcripts, is predominantly composed of IgG1 clones with 3 or more amino acid mutations, which is a signature of affinity maturation. All spike+ and RBD+ serum clones originating from IgG1 clones in the AB5 donor Ig repertoire are mature. The same trend is observed with serum clones in donors AB2 and AB6 (data not shown). The observation of BCR IgM clones as circulating IgG clones in serum shows the clones have undergone class-switching from IgM to IgG. Additionally, the observation that most IgG antibodies are mature, suggest that circulating antibodies matching the naive clones in the BCR repertoire are also somatically mutated.

**Figure 3.**
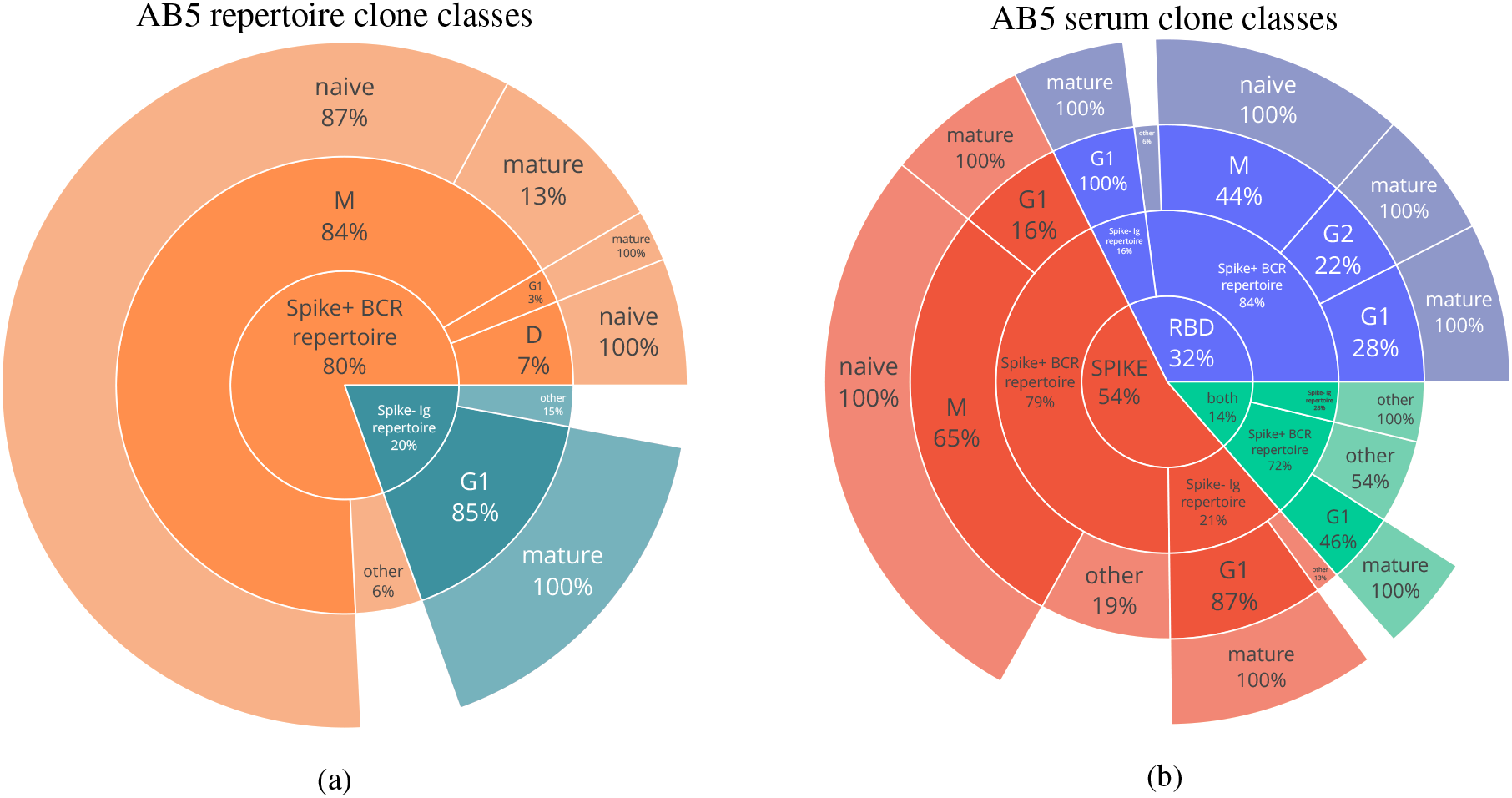
a) The class distribution of all sequences in the repertoire for donor AB5. The majority of antibodies in the BCR repertoire (orange) are IgM class with most sequences being naive (*<* 3 a.a. mutations), whereas the Ig repertoire (blue) are mature IgG class sequences. b) The antibody class distribution of serum clones (RBD in blue, spike in red, and both in green) shows enrichment of IgG class antibodies, but naive IgM class antibodies from the BCR repertoire are still predominant.

### 3.3 Characterization of serum antibodies show diverse and high-affinity binding to SARS-CoV2 domains and variants

Of the many antibodies observed in the BCR and Ig repertoires, antibodies circulating in serum are actively responding to infection. We selected 17 mature antibodies across the donors to express recombinantly as monoclonal antibodies (mAb) for further characterization. First, we tested reactivity in ELISA to Spike trimer, RBD, and NTD for the SARS-CoV-2 wild-type (WT) and Omicron subvariant B.1.1.529. To evaluate the potential of the antibodies to neutralize emerging variants, we further tested the antibodies in ELISA against the spike timer and RBD from Omicron subvariant BQ.1.1.

Two antibodies (5 and 12) from AB5 show binding to the spike protein, but not to any subdomain. Another two antibodies (4 and 7) from the same donor have binding to both the spike protein and the NTD across different Omicron variants (see Figure 4).

**Figure 4.**
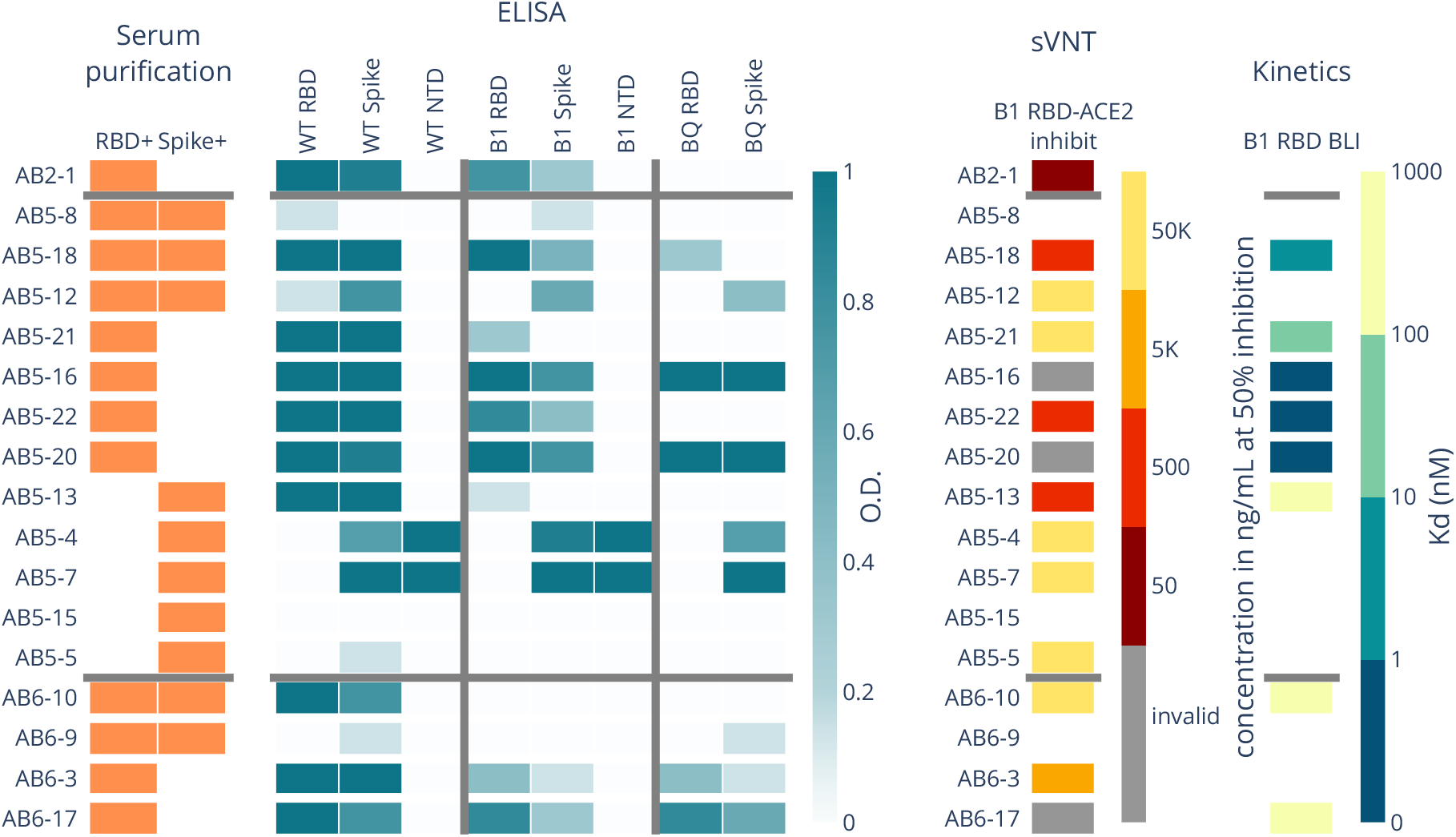
Serum RBD+ and spike+ clones from BCR repertoire were selected and recombinantly expressed and tested as mAbs. Panels show properties of each mAb. Serum purification panel shows which antigen purification the mAb is present in. ELISA panel shows binding activity as measured by optical density (O.D.) of each mAb to three SARS-CoV-2 variants and their RBD, spike, or NTD domains. Surrogate viral neutralization test (sVNT) shows concentration of mAb needed to inhibit 50% of B.1.1.529 RBD-ACE2 interactions. Kinetics panel shows the dissociation constant (Kd) of 8 mAbs tested (non-white) for B.1.1.529 RBD binding as measured by biolayer inteferometry (BLI).

Eleven antibodies show cross-reactivity to Omicron B.1.1.529, including antibody 1 from donor AB2 who had never been exposed to an Omicron variant of the virus. In addition, seven antibodies show cross-reactivity to Omicron subvariant BQ.1.1, which evolved after the peripheral blood was collected for this study.

We assayed the antibodies using surrogate viral neutralization testing (sVNT). Specifically, a competition ELISA was performed where B.1.1.529 RBD was immobilized, and each mAb was bound to the RBD at concentrations spanning 50,000 ng/mL to 5ng/mL. The complexes were then incubated with ACE2, and then washed to detect ACE2 binding to RBD. For each mAb, we tracked the lowest concentrations at which the mAb depleted 50% of the signal, which indicates that 50% of ACE2 was blocked from binding RBD. Six antibodies show potential for neutralization with 50% blocking at a concentration *<*5,000 ng/mL. Antibodies 16, 17, and 20 result in inconclusive inhibition values with increasing ACE2 binding at increasing mAb concentrations, but show no binding to ACE2 in ELISA (data not shown).

Strangely, these antibodies show binding activity to RBD and spike for all three SARS-CoV2 variants, with strong affinity to B.1.1.529 RBD.

For antibodies with ELISA activity to RBD from donors AB5 and AB6, we measured the binding kinetics to B.1.1.529 RBD using biolayer inteferometry. Antibodies 10, 13, and 17 have dissociation constants (Kd) ranging from 171nM to 723nM, and antibodies 18 and 21 had low Kd of 3.4nM and 30nM, respectively. Antibodies 16, 20, and 22 showed no dissociation from RBD, and are likely sub-nanomolar affinity antibodies that are below the detection limits of the instrument.

### 3.4 Serum antibodies showed similarity to previously known public clones

Clones that are highly similar across donors, in terms of originating heavy chain germline V-gene and CDR3 sequence, suggest convergent immune responses, and are referred to as public clones. Public clones to SARS-CoV-2 have been previously reported [19], and known anti-SARS-CoV-2 antibodies have been collected in the coronavirus antibody database (CoV-AbDab). We compared our Ig and BCR repertoires, and serum selected antibodies to previously reported sequences, and found characterization data provided in CoV-AbDab to agree with that of similar mAbs in our list of 17. For example, AB5-18, which binds to RBD and neutralizes B.1.1.529 matches antibody BD56-411 with 83% HCDR3 similarity and 91% HC similarity (Figure 5). BD56-411 is reported to bind RBD Omicron, including neutralization of subvariants B.1.1.529 to BA.2. All the serum clones with public clone matches originate from the BCR repertoire. Additionally, 76, 95, and 38 public clones in the BCR and Ig repertoires for donors AB2, AB5, and AB6, respectively, were not observed in serum. This provides alternate confirmation that the serological compartment retains a narrow population of antibodies that are distinct from the cellular repertoire.

**Figure 5.**
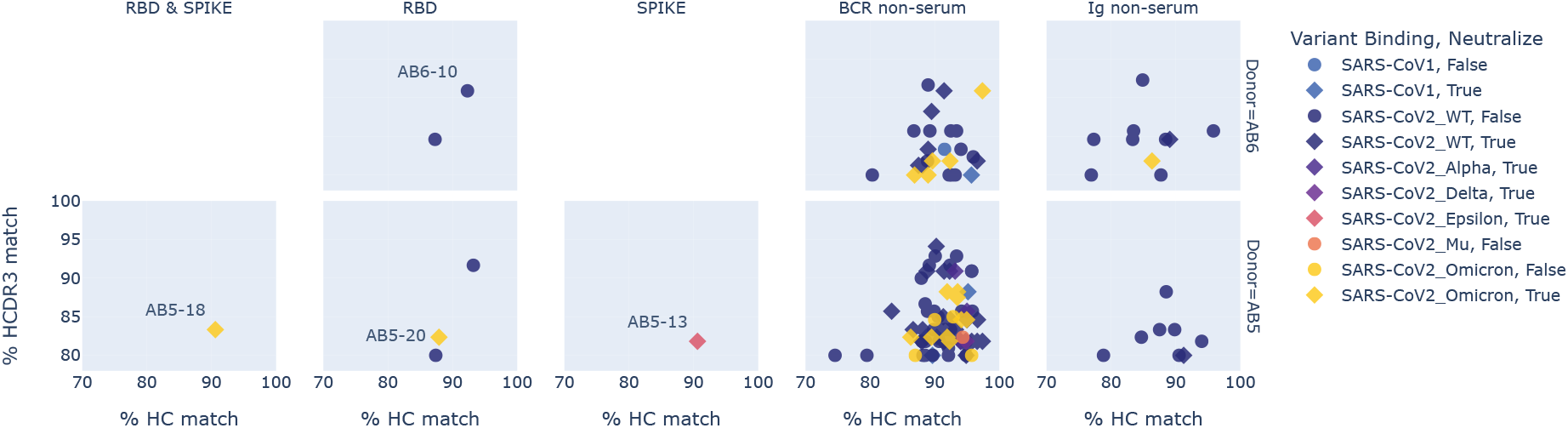
Propagation of activity profiles from public CoV-AbDab sequences to sequences observed in donors AB5 and AB6. Each column shows a source of the donor-derived sequence, observed in serum binding to RBD & SPIKE, RBD-only, SPIKE-only, or present in BCR or Ig repertoire. The similarity of each donor-derived sequence is shown by % HCDR3 match on the y-axis, and % HC match on the x-axis to a CoV-AbDab sequence. Markers are colored by the most recent variant binding, and styled as neutralizating (diamond) or non-neutralizing (circle) as reported in Cov-AbDab. Recombinant mAbs 10, 13, 18, and 20 matched public clones, and our characterization data showed the same activity profiles as reported public sequences.

## 4 Discussion

Of the more than 100,000 clones in the BCR and Ig repertoire across the three donors in this study, less than *<*0.1% are represented as anti-SARS-CoV-2 Omicron variant antibodies. The B-cell repertoire in peripheral blood is a weak proxy for representing the dynamics of active antibodies circulating in serum, as shown by other proteogenomic studies [13, 10, 24]. In serum, 70%-80% of secreted antibodies are IgG class by mass. Conversely, the cellular repertoire accessible in peripheral blood is primarily naive-mature B cells, and memory B cells expressing surface IgM.

Plasma cells and plasmablasts, which secrete antibody, account for *<*0.5% of lymphocytes in peripheral blood, and 60% express IgA class antibodies. Predictably, a large proportion of serum-observed IgG clones are of class IgM in the BCR repertoire (46%-76% between donors), which suggests that these clone lineages are undergoing expansion, and are present in donors with multiple cell types.

Peripheral blood was collected from each donor at least four months after their self-reported exposure to viral antigens through vaccination or infection to uncover a significant proportion of B cells and circulating antibodies still responding to SARS-CoV-2 antigens with our Alicanto platform. Two donors were exposed to an early Omicron variant by breakthrough infection, and had significantly more spike+ (Omicron variant B.1.1.529) B cells, and circulating antibody matching RBD+ and spike+ clones than the donor who was only exposed to SARS-CoV-2 wild-type antigens via hybrid immunity–infection followed by vaccination. Previous proteogenomic studies [26, 28] had reported that imprinting effects of natural SARS-CoV-2 infection versus vaccination directs immune response to divergent epitopes, where vaccinated individuals have more circulating clones directed to RBD than non-RBD spike proteins. We did not find a compelling trend in our data to support the previous finding. The donor with initial exposure by natural infection had only slightly more spike+ serum clones than RBD+, and donors who were vaccinated prior to infection had either slightly more RBD+ serum clones than spike+ clones, or vice versa. One rationale for the difference could be the dynamics of serum antibodies, and proportion of epitope focus may change depending on recent versus longitudinally distant infection.

Using our Alicanto platform, we selected highly confident serum clones observed in the BCR repertoire to recombinantly express for further validation and characterization. Of the candidates selected, we found diverse binding modalities to the RBD, NTD, and non-RBD/NTD of the spike protein. Candidates from donors exposed to an early Omicron variant were observed to be cross-reactive to the wild-type strain and other Omicron variants, including BQ.1.1, an escape variant discovered many months later. Two candidates from a donor with breakthrough infection bound to the tested RBD Omicron variants, but dissociation kinetics were too slow to be reliably measured by BLI. These candidates are likely strongly neutralizing with sub-nanomolar dissociation constants.

To further characterize clones observed in the BCR, Ig, and serum repertoires, we leverage sequences and annotations in CoV-SAbDab [19] to investigate convergent immune responses. Across the different donors, 209 clones match to publicly reported sequences, i.e. public clones, with seven clones present in serum. Four of the serum clones had been recombinantly expressed, and have strong agreement with neutralization activity and variant binding to the reported public clones. Public clones are also present in the BCR repertoire, corroborating evidence that the antigen-specific antibodies circulating in serum represent a distinct dynamic from the antigen-specific B cell repertoires.

Previous findings of SARS-CoV-2 reactive B cells have primarily been IgM class with few somatic mutations [12]. Likewise, our spike+ BCR repertoires have fewer somatic mutations compared to the Ig repertoire. Circulating IgG in serum is a product of affinity maturation, and clones that match the Ig repertoire are all somatically mutated in the repertoire (*>*3 amino acid mutations). In contrast, most of the clones matching the BCR repertoire are naive. This suggests that the circulating antibodies in serum that match the BCR repertoire likely contain missed somatic mutations, and a potential boon for naturally evolved and higher affinity anti-SARS-CoV-2 antibodies. A limitation of the proteogenomic method is that the serum clone needs representation in the BCR repertoire to be observed and tested as a recombinant monoclonal. A promising direction is to directly sequence antibodies *de novo* without a repertoire. Existing efforts have involved a significant amount of sample preparation and mass spectrometry instrument time and labor, but show potential for deriving a functioning antibody sequence from IgG protein alone is possible [11, 1]. Continual development of mass spectrometry and informatic methods is necessary for an in-depth characterization of the multitude of antibodies circulating in serum.

## 5 Acknowledgments

We thank Miin Lin for contributions in reviewing and editing the manuscript.

